# Automated Pipeline for Comparing Protein Conformational States in the PDB to AlphaFold2 Predictions

**DOI:** 10.1101/2023.07.13.545008

**Authors:** Joseph I. J. Ellaway, Stephen Anyango, Sreenath Nair, Hossam A. Zaki, Nurul Nadzirin, Harold R. Powell, Aleksandras Gutmanas, Mihaly Varadi, Sameer Velankar

## Abstract

Proteins, as molecular machines, are necessarily dynamic macromolecules that carry out essential cellular functions. Recognising their stable conformations is important for understanding the molecular mechanisms of disease. While AI-based computational methods have enabled protein structure prediction, the prediction of protein dynamics remains a challenge. Here, we present a deterministic pipeline that clusters experimentally determined protein structures to comprehensively recognise conformational states across the Protein Data Bank. Our approach clusters protein chains based on a GLObal CONformation (GLOCON) difference score, which is computed from pairwise C-alpha distances. By superposing the clustered structures, differences and similarities in conformational states can be observed. Additionally, we offer users the ability to superpose predicted models from the AlphaFold Database to the clusters of PDB structures. This clustering pipeline significantly advances researchers’ ability to explore the conformational landscape within the PDB. All clustered and superposed models can be viewed in Mol* on the PDBe Knowledge Base website, or accessed in as raw annotations via our GraphAPI and FTP server. The clustering package is made available as an open-source Python3 package under the Apache-2.0 license.

## Main

Proteins are molecular machines that drive biological processes, adopting specific shapes – or *conformations* – to carry out cellular functions. A detailed understanding of protein dynamics is essential for elucidating the molecular mechanisms of diseases, such as SARS-CoV2 infection or cancer^1,2^. The structural states of proteins can be influenced by a range of factors, including experimental technique, ligand presence, oligomeric state, and experimental conditions^3^. In recent years, the emergence of AI-based computational methods for predicting protein structures has enabled access to protein structure models lacking experimentally determined structures^4^. However, they are yet to provide information on protein dynamics. Therefore, analysis of protein dynamics in this era of data abundance requires a multi-faceted approach, integrating both experimental and computational techniques^5,6^. Such an approach would provide a more comprehensive picture of the ever-expanding universe of known protein structures, and assist the development of novel therapies and drugs modulating protein dynamics^7^.

The Protein Data Bank (PDB) provides a rich sampling of protein conformational space, where many sequences are represented by independently solved structures. Much of the biologically meaningful conformational space is captured, but model redundancy – revealing structural variability – is yet to be fully exploited^3,8^. At present, the PDB offers only the ‘title’ field to describe conformational states, but information is often intermittently provided or not known. For example, adenylate kinase from *Escherichia coli* (UniProt accession: P69441) adopts either an *open* or *closed* conformation^9^, but many PDB entries ignore conformational state in their title.

Even where descriptions of protein dynamics are included, the information is typically insufficient to highlight structural similarities or differences between peptides in the same or different conformations. For example, the individual PDB entries of liganded-bound human aldose reductase (UniProt accession: P15121) fail to collectively capture the structural heterogeneity in the β-sheet region spanning Val121-Arg156. Additionally, new insights could be made when aggregating known structures, such as the closeness in structure between MurD’s *open* and *closed* conformations relative to its *intermediate* state (UniProt accession: P14900).

These limitations pose a significant challenge for programmatically assessing the dynamics of a single protein, let alone the entire PDB. Structured labels, not free-text descriptions, are needed for systematic analysis of the structural diversity across the archive – enabling exploration of the conformational space contained in the PDB.

To address these challenges, the PDBe has developed a deterministic clustering pipeline for protein conformational states that superposes and clusters protein chains across the PDB archive. The pipeline aims to capture structural heterogeneity and enable accurate and automated identification of distinct conformational states. Our open-source pipeline updates data weekly and is accessible through the PDBe-KB^10^.

In outline, peptides from the PDB with 100% sequence identity are binned into groups called *segments* (**Fig. 1a**) and superposed. Pairwise Ca distances for all peptides mapped to a segment are measured and used to compute a GLObal CONformation difference score (*GLOCON score*) for each chain-chain comparison, adjusted to penalised gaps (**Fig. 1b**) **[Supplementary Materials]**. UPGMA agglomerative clustering is finally performed to cluster chains based on the GLOCON scores, splitting the segment into *clusters* – recognised conformational states (**Fig. 1c-d**).

**Figure 1.**
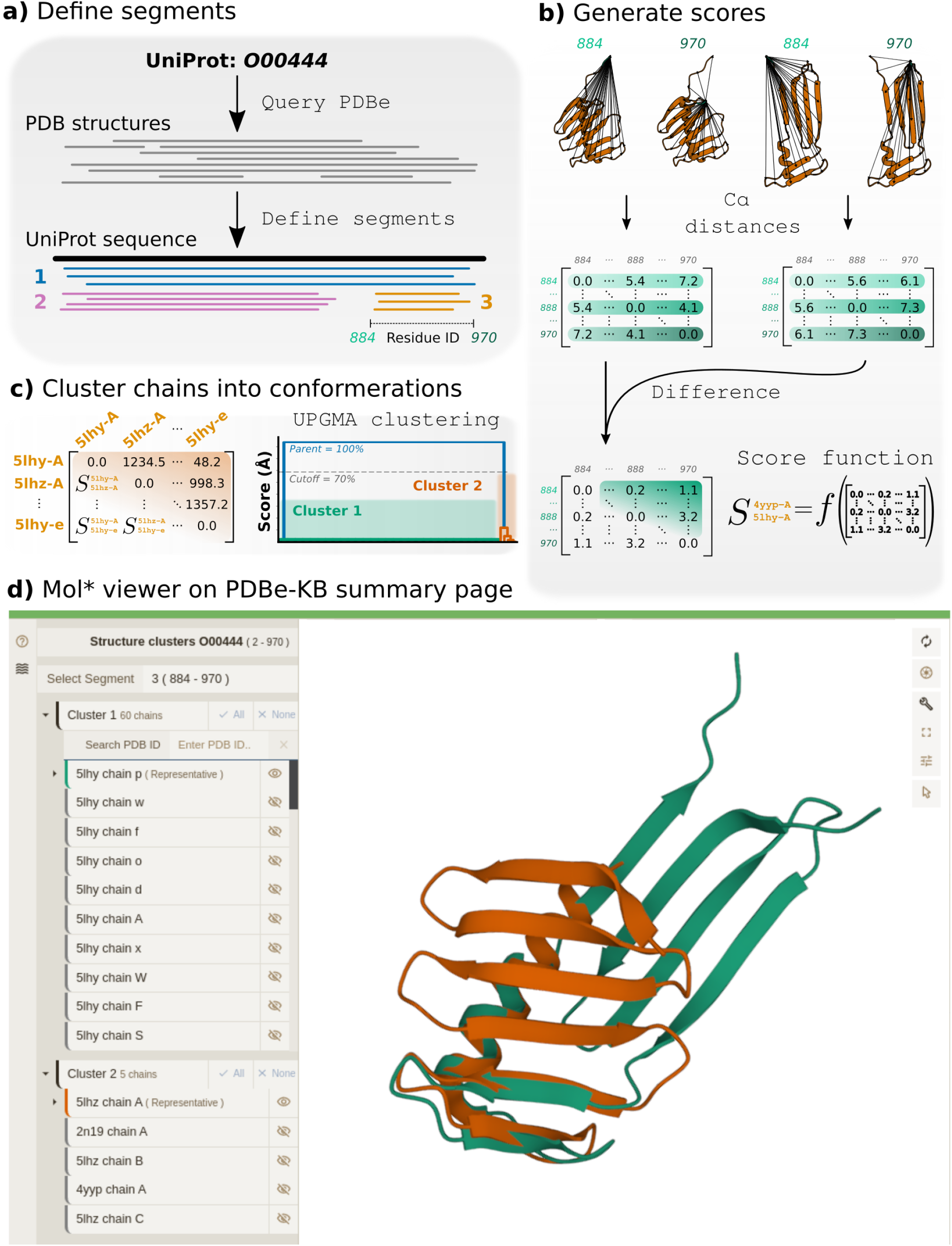
Recognition of protein conformational states across the PDB archive Outline of the conformational state recognition pipeline, run weekly on the PDB archive and made available through the PDBe-KB. **a)** All chains pertaining to a given UniProt accession (100% sequence identity) are assigned to segments, based on their overlap with the reference UniProt sequence. Non-overlapping sequences are grouped into separate segments. **b)** Chains are superposed to all other chains within their assigned segment. **c)** Chain-chainGLOCON scores are calculated for all peptides within a segment, before **d)** agglomerative clustering is performed. The results are displayed on the PDBe-KB’s Summary page, using the Mol* viewing window.

We offer users the ability to superpose predicted models onto the PDB structural clusters by taking advantage of the millions of high-accuracy protein structure predictions in the AlphaFold Protein Structure Database (AFDB). The root-mean-square deviation of the AlphaFold2 (AF2) model from each cluster’s representative chain is calculated and displayed, allowing identification of the conformational state predicted by the AF2 pipeline. By default, we blur disordered regions of the AF2 models that have low confidence (i.e. pLDDT) scores. This comparison allows users to quickly identify which conformational state AlphaFold predicts, potentially expediting functional determination.

In conclusion, the clustering pipeline presented here significantly advances researchers’ ability to interrogate the conformational landscape within the PDB and compare it to AFDB models. By delivering structured conformational state annotations, using this systematic approach, accurate identification of structural variation is now possible. Biologists can quickly interpret conformational diversity via a browser or access archive-wide information programmatically. Moreover, predicting conformational states represents a starting point for automating the assignment of biological context to protein dynamics. Researchers worldwide now have access to an invaluable tool for studying protein conformations and advancing our understanding of molecular mechanisms in health and disease.

## Data availability

All clustering results for the PDB archive (covering over 60k+ UniProt accessions) are updated weekly and can be accessed directly from the PDBe-KB’s graph API (e.g. https://www.ebi.ac.uk/pdbe/graph-api/uniprot/superposition/<uniprot_accession >). Dendrograms of clustering results and per-chain rotation translation matrices are also available via our FTP server. A manually-curated benchmark was compiled for testing code; all the data are distributed under the CCBY-4.0 license.

## Code availability

The conformer prediction code is written in Python3.10 and is available as a pip-installable package (`pip install protein-cluster-conformers`). Users can cluster chains directly on their local machine using the available Python wrappers in the GitHub repository.

## Acknowledgements

The authors would like to thank the UKRI-Biotechnology and Biological Sciences Research Council for providing funding under the FunCLAN (BB/V016113/1) project and the European Molecular Biology Laboratory-European Bioinformatics Insititute for supporting development of the service.

## Notes

https://github.com/PDBeurope/protein-cluster-conformers

https://www.kaggle.com/datasets/josephellaway/distinct-monomeric-protein-conformers

https://ftp.ebi.ac.uk/pub/databases/pdbe-kb/benchmarking/distinct-monomer-conformers/

### Competing Interest Statement

The authors have declared no competing interest.

### Summary of Updates

Name correction

