## Supplementary information for "Automated Pipeline for Comparing Protein Conformational States in the PDB to AlphaFold2 Predictions"

### Identifying Protein Conformational States in the PDB and Comparison to AlphaFold2 Predictions:

July 11, 2023

1. European Bioinformatics Institute, Protein Data Bank in Europe, Hinxton, UK
2. The Warren Alpert Medical School of Brown University, Providence, RI, USA
3. Imperial College London, Department of Life Sciences, London, UK
4. WaveBreak Therapeutics Ltd., Clarendon House, Clarendon Road, Cambridge, UK

**2 Examples of conformational state recognition** 6

**3 Data and Code Availability** 9

**4 References** 9

### 1 Supplementary Methods

#### 1.1 Segment definition

The pipeline is initially parsed a single UniProt ID, from which it queries the PDB archive [1–3] to identify all peptide structures overlapping any region of the given UniProt [4] sequence. These retrieved structures are next separated into '*segments*', or collections of peptide sequences with 100% sequence identity (ignoring non-modelled residues and point mutations), to be grouped based on their sequence overlap with each other. These *segments* are determined by calculating a pair-wise dissimilarity matrix of overlaps between the sequences of all retrieved structures. The elements in the dissimilarity matrix are reverse Jaccard scores (1-Jaccard) between all chains pertaining to the given UniProt (Eq. 1).

$$J_{mn} = 1 - \frac{|\mathbf{R_m} \cap \mathbf{R_n}|}{|\mathbf{R_m} \cup \mathbf{R_n}|} \tag{1}$$

where $\mathbf{R_m}$ and $\mathbf{R_n}$ are the sets of SIFTS-allocated [5] UniProt residue IDs (integers) of chains $m$ and $n$ in the segment. These sets can be formally described as (Eq. 2):

$$\mathbf{R_m} = \{1, 2, \ldots, r_m\}, \qquad \mathbf{R_n} = \{1, 2, \ldots, r_n\}, \qquad \forall \;\; r_{m,n} \in \mathbb{Z}^+ \tag{2}$$

$r_m$ and $r_n$ denote the C-terminal most residue IDs, and their sets $\mathbf{R_m}$ and $\mathbf{R_n}$ need not be continuous integer arrays, capturing any true occurrences of residues being omitted from the parsed structure. PDB entries with complete sequence overlap or none obtain reverse Jaccard scores of 0 and 1, respectively. Sequences below 20 residues are ignored. A hierarchy is constructed based on the dissimilarity matrix and cutoff at 0.99 to define segments. Branch points in the hierarchy below this cutoff are assigned to a segment. Although not yet observed in the archive, this process could result in as many segments as retrieved chains. In practice, the number of segments is typically in the range of 1-4.

For instructional purposes, let us denote one segment (of potentially many) by $\mathbf{P}$, which naturally defines the set of all chains in the segment. Chains $m$ and $n$ in segment set $\mathbf{P}$ can be represented as integers within the set $\mathbf{P}$:

$$\mathbf{P} = \{m, \ldots, n\}, \qquad \forall \;\; m, n \in \mathbb{Z}^+ \tag{3}$$

This establishes the notation used to describe clustering throughout the rest of this section.

#### 1.2 C$\alpha$-distance function

Regardless of the original structure's oligomeric state, all chains within segment $\mathbf{P}$ are clustered based on a pair-wise C$\alpha$-distance difference score. The score is derived by first generating an all-to-all intra-sequence C$\alpha$ distance matrix, within each polypeptide chain, for all chains in the segment. Eq.4 illustrates the generation of such internal C$\alpha$-distance matrices for chains in $\mathbf{P}$.

$$\mathbf{C_1} = \begin{bmatrix} c^1_{11} & c^1_{12} & \cdots & c^1_{1i} \\ c^1_{21} & c^1_{22} & \cdots & c^1_{2i} \\ \vdots & \vdots & \ddots & \vdots \\ c^1_{j1} & c^1_{j2} & \cdots & c^1_{ij} \end{bmatrix}, \; \mathbf{C_2} = \begin{bmatrix} c^2_{11} & c^2_{12} & \cdots & c^2_{1i} \\ c^2_{21} & c^2_{22} & \cdots & c^2_{2i} \\ \vdots & \vdots & \ddots & \vdots \\ c^2_{j1} & c^2_{j2} & \cdots & c^2_{ij} \end{bmatrix}, \quad \cdots, \; \mathbf{C_m} = \begin{bmatrix} c^m_{11} & c^m_{12} & \cdots & c^m_{1i} \\ c^m_{21} & c^m_{22} & \cdots & c^m_{2i} \\ \vdots & \vdots & \ddots & \vdots \\ c^m_{j1} & c^m_{j2} & \cdots & c^m_{ij} \end{bmatrix} \tag{4}$$

$\forall \;\; i \in \mathbf{R_m}$ where $i_{max} = max\left(\mathbf{R_m}\right) = r_m$ and $m \in \mathbf{P}$. Individual elements represent the Euclidean C$\alpha$-C$\alpha$ distance between all residues in the chain.

$$c^m_{ij} = d\left(C\alpha_i, C\alpha_j\right) \quad \in \mathbb{R}^+ \tag{5}$$

$$= \sqrt{\left(C\alpha^x_j - C\alpha^x_i\right)^2 + \left(C\alpha^y_j - C\alpha^y_i\right)^2 + \left(C\alpha^z_j - C\alpha^z_i\right)^2} \tag{6}$$

Therefore, $\mathbf{C_m}$ denotes the general expression used to capture all internal C$\alpha$ distances in chain $m$ for the segment $\mathbf{P}$. We can further simply the expression for $C_m$ in Eq. 4 as:

$$\mathbf{C_m} = \left[c^m_{ij}\right] \;\; \forall \;\; i, j \in \mathbf{R_m} \tag{7}$$

In total, the number of $\mathbf{C_m}$ matrices to compute equals the size of set $\mathbf{P}$, or $|\mathbf{P}|$, and can be denoted by the set $\mathbf{Q}$, where:

$$\mathbf{Q} = \{\mathbf{C_1}, \mathbf{C_2}, \cdots, \mathbf{C_m}\} \tag{8}$$

$$|\mathbf{Q}| = |\mathbf{P}| \tag{9}$$

##### 1.3 Distance-difference function

The global change in backbone C$\alpha$ positions between all chains in segment $\mathbf{P}$ is captured by computing the absolute difference between paired elements across all matrices contained in $\mathbf{Q}$. Parity between elements (C$\alpha$-C$\alpha$ Euclidean distances) in the matrices contained in $\mathbf{Q}$ is established using the UniProt residue indexing performed by the PDBe's SIFTs process [6, 7].

$$\mathbf{D_{mn}} = abs\left(\mathbf{C_m} - \mathbf{C_n}\right) \tag{10}$$

$$= \begin{bmatrix} d_{11}^{mn} & d_{12}^{mn} & \cdots & d_{1j}^{mn} \\ d_{21}^{mn} & d_{22}^{mn} & \cdots & d_{2j}^{mn} \\ \vdots & \vdots & \ddots & \vdots \\ d_{i1}^{mn} & d_{i2}^{mn} & \cdots & d_{ij}^{mn} \end{bmatrix} \tag{11}$$

$$= \left[d_{ij}^{mn}\right] \quad \forall \quad i, j \in \mathbf{R_m} \cap \mathbf{R_n} \tag{12}$$

where $d_{ij}^{mn} = |c_{ij}^{m} - c_{ij}^{n}| \quad \forall \ d_{ij}^{mn} \in \mathbb{R}$. Intuitively, we should expect $\mathbf{D_{mn}} \equiv \mathbf{D_{nm}} \quad \forall \quad m, n \in \mathbf{P}$ and $\mathbf{D_{mn}} = \mathbf{D_{nm}} = 0_{ij}$ (empty matrix) where $n = m$. A total of $|\mathbf{Q}|^2$ distance difference matrices ($\mathbf{D_{mn}}$) are required to capture all chain-chain C$\alpha$ comparisons in the segment. In practice, only one triangle (excluding the diagonal for $n = m$) contains unique C$\alpha$ distance differences; Fig.1 illustrates the observed differences in the distance-difference matrices between chains within and between conformational states. Plotting the C$\alpha$ distance difference matrices like this shows there is clearly a large C-terminal domain shift in the backbone, from ˜residue 250, suggesting a conformational state difference between stated chains.

##### 1.4 Score function

Once distance difference matrices ($\mathbf{D_{mn}}$) are computed, we apply a 3 Å cutoff to all elements in the array [8]. Values below this cutoff are set to zero according to Eq.14:

$$d_{ij}^{mn}* = \begin{cases} d_{ij}^{mn}, & \text{if } d_{ij}^{mn} \geq 3 \\ 0, & \text{else} \end{cases} \tag{13}$$

$$\mathbf{D_{mn}^*} = \left[d_{ij}^{mn}*\right] \quad \forall \quad i, j \in \mathbf{R_m} \cap \mathbf{R_n} \tag{14}$$

This acts as a filter to prevent small differences from influencing the final score [8]. In outline, the score is simply the sum of the upper triangle of the distance difference matrices between all chain-chain comparisons within the segment $\mathbf{P}$. To penalise sequence gaps and partial overlaps at the termini, we normalise the sum of the upper triangle to the fraction of common residues between each chain over the product of the

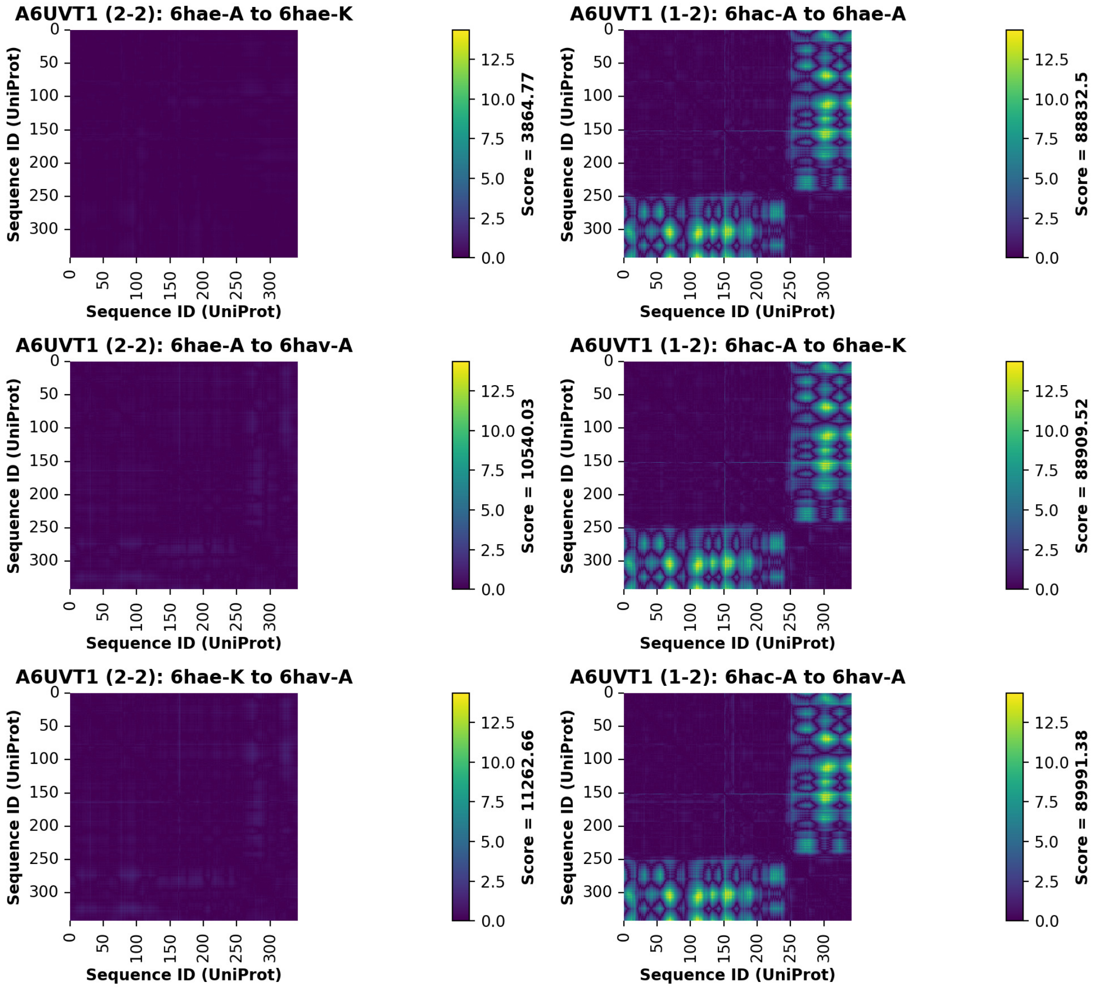

Figure 1: **Example of expected distance-difference matrices for chains within (*left*) and between (*right*) conformations**. The 5,10-methenyltetrahydromethanopterin hydrogenase enzyme from *Methanococcus aeolicus* (UniProt accession: A6UVT1) is chosen due to one structure in the segment exhibiting a well-defined C-terminal domain movement. This can be seen by the green-yellow regions of the plots of chains compared in different conformations. GLOCON scores (in Å) are provided to the right of the colour bars.

length of the two sequences. A scaler between 0-1 is returned, denoting zero and complete sequence overlap respectively. Eq.15 describes this score in full for all common residues between chains $m$ and $n$.

$$S_{mn} = \frac{|\mathbf{R_m} \cap \mathbf{R_n}|^2}{|\mathbf{R_m}| \cdot |\mathbf{R_n}|} \cdot \sum_{ij}^{i<j} \left[ d_{ij}^{mn} * \right] \quad \forall \quad i, j \in \mathbf{R_m} \cap \mathbf{R_n} \tag{15}$$

**GLOCON score**

where $S_{mn} \equiv S_{nm} \;\; \forall \;\; m, n \in \mathbf{P}$. A total of $\frac{|\mathbf{P}| \cdot (|\mathbf{P}|-1)}{2}$ scores, $S_{mn}$ are computed, amounting to all unique chain-chain comparisons within the segment, excluding $S_{mn}$ for $m = n$.

#### 1.5 Clustering algorithm

Using the score function from Eq.15, a dissimilarity matrix is compiled for all the segment set, $\mathbf{P}$:

$$\mathbf{S} = \left[S_{mn}\right] \quad \forall \quad m, n \in \mathbf{P} \tag{16}$$

This dissimilarity matrix is used for UPGMA agglomerative clustering, grouping chain pairs with the smallest scores to build up a hierarchical tree [9]. A threshold of 70 % of the parent node's score is applied, grouping all nodes below this cutoff into recognised conformers, or *clusters* (Fig.2). These clusters are therefore determined solely from global backbone C$\alpha$-C$\alpha$ differences, effectively decoupling conformational state prediction from structural alignment.

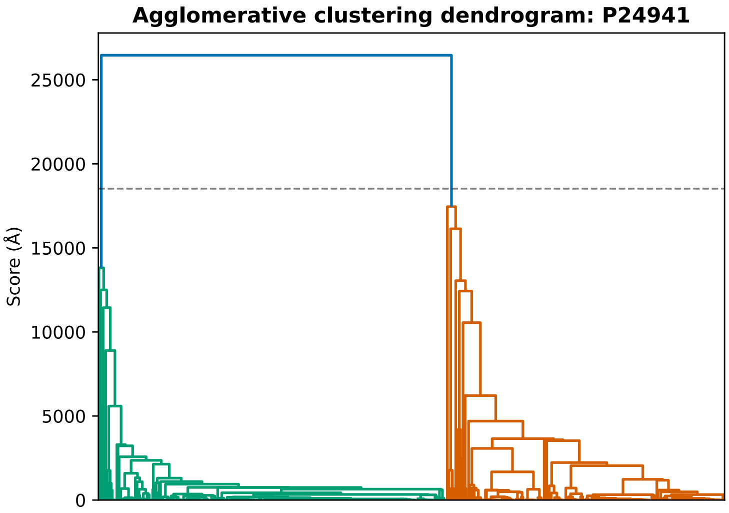

Figure 2: **Example of conformational state recognition: Agglomerative clustering results for CDK2 in dendrogram form**. The y-axis denotes the average score computed during grouping via UPGMA clustering. Labels along the x-axis have been omitted for clarity. In this case, two conformations have been recognised.

Cluster information is updated weekly to accommodate newly solved PDB entries or modifications to existing structures. Checks are in place to avoid matrix recalculation and parallelism is employed to distribute linear algebra operations across available processors. We recommend using ten processes for optimal runtime on an SSD storage device, but running on a single thread is completely feasible.

#### 1.6 Structural alignment and Mol* viewer

Once clusters have been defined, we *superpose* (structurally align) all chains within segment **P** using the GESAMT aligner available from the CCP4 suite [5]. GESAMT uses the Q-score metric – a modified version of RMSD – to perform superposition, and obtains excellent alignment around structurally conserved regions due to its chain fragmentation approach, whilst allowing the rest of the molecule to deviate. This can better highlight regions of structural dissimilarity as conserved regions are favourable to superposition by GESAMT. Occasionally, GESAMT fails to align chains (especially where non-terminal spans of residues are not modelled), in which case the secondary structure-based aligner SSM is deployed instead [10]. These structural alignment results are loaded into the PDBe-KB's Aggregate Views of Proteins summary page (Fig.3) [3]. Representatives for each cluster are determined based on model quality and sequence length.

#### 1.7 Integration with AlphaFold database

PDBe-KB is now more tightly integrated with the AlphaFold database [3, 11]; a structure's corresponding AlphaFold model automatically superposes to the representative chains from each cluster when the "Load AlphaFold structure" button is clicked. Structural alignment is performed on the client's machine, although the PAE matrix and pLDDTs are pre-generated. Deep blue denotes high pLDDT regions and a transparency

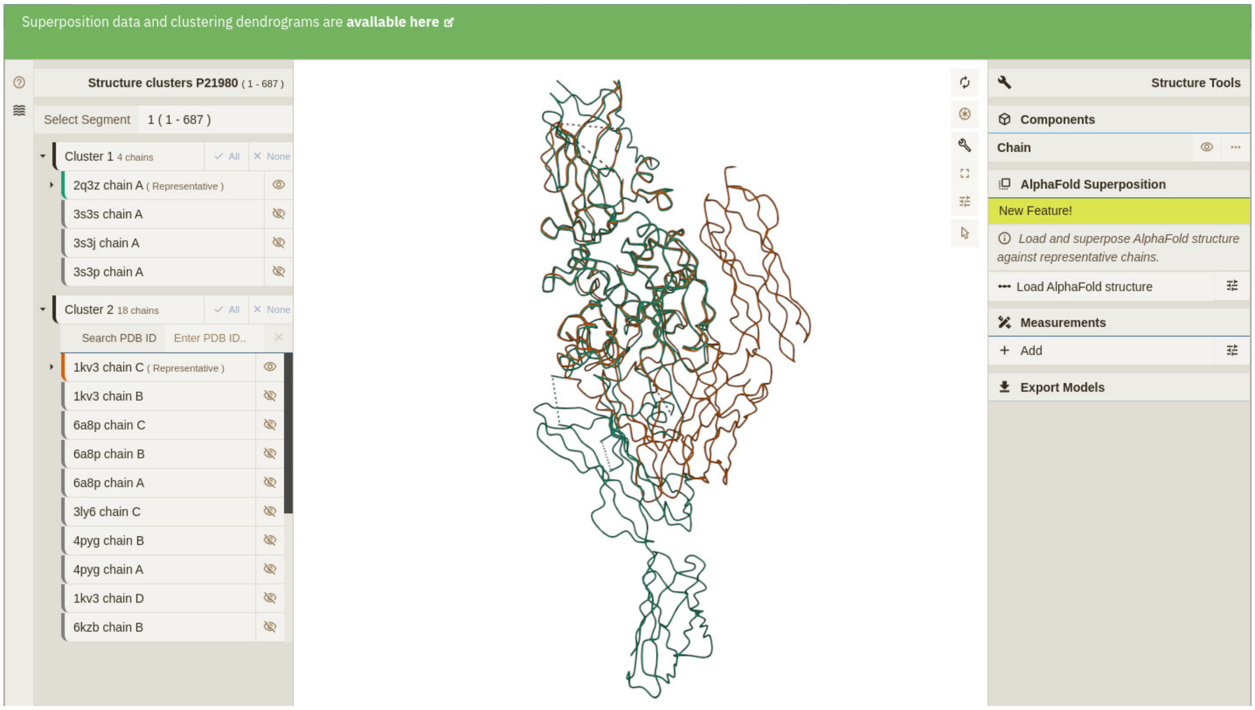

Figure 3: **Recognised conformational states displayed as a superposed ensemble in PDBe-KB's Mol* viewer.** Up-to-date documentation on the data displayed by this window can be found on the PDBe-KB's GitHub manual repository By default, only representative chains are displayed for performance. 'Clusters' named here are the recognised conformational states predicted by the UPGMA algorithm in §1.5. Individual chains within any cluster can be revealed by checking the eye icon right of the PDB-chain code. Users can also search for chains within a cluster if they contain many structures. Segments (if multiple are available) can be navigated using the "Select Segments" dropdown in the top left of the window. Dendrograms of cluster assignment and rotation-translation matrices can be accessed via the link in the green window header.

threshold can be toggled in the right-hand panel. The RMSD for the superposed AlphaFold model to each cluster's representative chain is displayed in the right-hand panel, along

If running the code locally, you can also include the AlphaFold structure in your recognition of conformational states. The AlphaFold structure will be included in the resulting dendrogram, getting assigned to a cluster along with your other parsed structures. In order to deliver our integrated AlphaFold view, we do not currently run this feature on a weekly basis; RMSD could, however, be used to infer conformational assignment.

#### 2 Examples of conformational state recognition

Before September 2022, Q-score (used to assess structural alignments by GESAMT) score was used for UPGMA clustering. Although often successful for structures where the entire peptide was considered for the computation of Q-score, the metric incorrectly under- and over-clustered many UniProt segments. This prompted us to compile and manually curate a small dataset of conformationally distinct monomeric proteins. The dataset was then used to check if the new score could better separate protein chains by structural dissimilarity.

Shown in Fig.5 are a selection of examples in which the GLOCON score-based clustering pipeline is able to recognise the biologically meaningful conformational states successfully. For example, hexokinase from *Sulfurisphaera tokodaii* (UniProt accession: Q96Y14) – the first enzyme in the glycolytic pathway – is responsible for initialising the process of respiration and highly important in anaerobic conditions. The

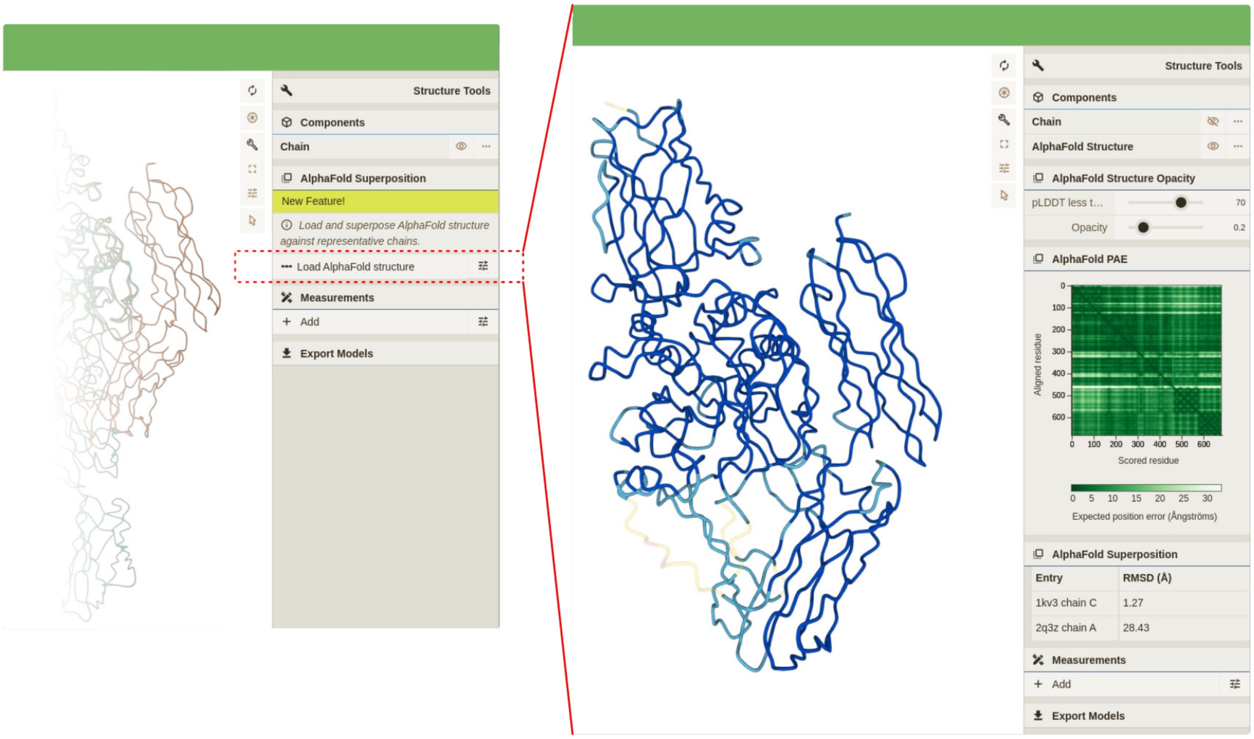

Figure 4: **Illustration of the AlphaFold structure imported into the PDBe-KB's Mol* viewer**. Left-hand window shows the window before the "Load AlphaFold structure" button is clicked, and the right-hand side window shows the rendered AlphaFold structure. In the right-hand window, additional information on the AlphaFold model is displayed, including RMSD to each cluster's representative chain.

kinase is moderately promiscuous to sugar substrates, as mentioned by [12]. It can associate with glucose, mannose, glucoseamine, xylose and N-acetylglucosamine (and possibly more). As these authors identify, hexokinase adopts an open or closed conformation, dependent on sugar binding, although ADP binding has marginal effect on the shape of the protein. Encouragingly, we are able to discern between these open and closed states and (due to our pipeline working on a per-chain basis) able to separate the open and closed chains solved within the asymmetric unit of 2E2Q (Fig.5a).

Additionally, human aldose reductase (UniProt accession: P15121) accepts a diverse range of carbonyl-base substrates, reducing them to alcohol products using NADH as an electron source. It has impressive structural redundancy in the ligand-bound form. In other words, many structures of the protein have been independently solved with a multitude of ligands, providing a wealth of information on the conformational heterogeneity within this ligand conformation [13]. We are able to separate the only non-liganded structure in the PDB (1XGD) from all other ligand-bound chains. Separation from and superposition with all other chains highlights that the unliganded structure possesses a large deviation in the P211-232 loop region, which is characterised far less than the liganded structures. The new tool clearly exposes this difference and also facilitates immediate interpretation of the structural differences between liganded structures (Fig.5b).

Furthermore, the circadian rhythm protein KaiB helps regulate the day-night cycle in cyanobacteria. Associating with KaiA and KaiC, KaiB from *Thermosynechococcus vestitus* partakes in a concerted cycle of complex formation, autophosphorylation and autodephosphorylation of KaiC, completing each oscillation every $\sim$ 24 hours [14]. Because we only cluster single protein chains, we are able to identify the ground and fold-switch states. One of the KaiABC oscillatory complex components regulating the day-night circadian clock in cyanobacteria, KaiB adopts a homotetrameric ground state (Fig.5c, left/teal) during the day and a thioredoxin-like "fold-switch" state at night (Fig.5c, right/orange). The fold-switch state is ordinarily stabilised upon oligomerisation with KaiC and KaiB subunits, forming a large multimeric complex [14]. The clustering method described herein identifies the structures solved in these two states, and shows the protein's AlphaFold model (Fig.5c, blue) is closer in conformation to the night-dominant fold-switch state (Fig.5c).

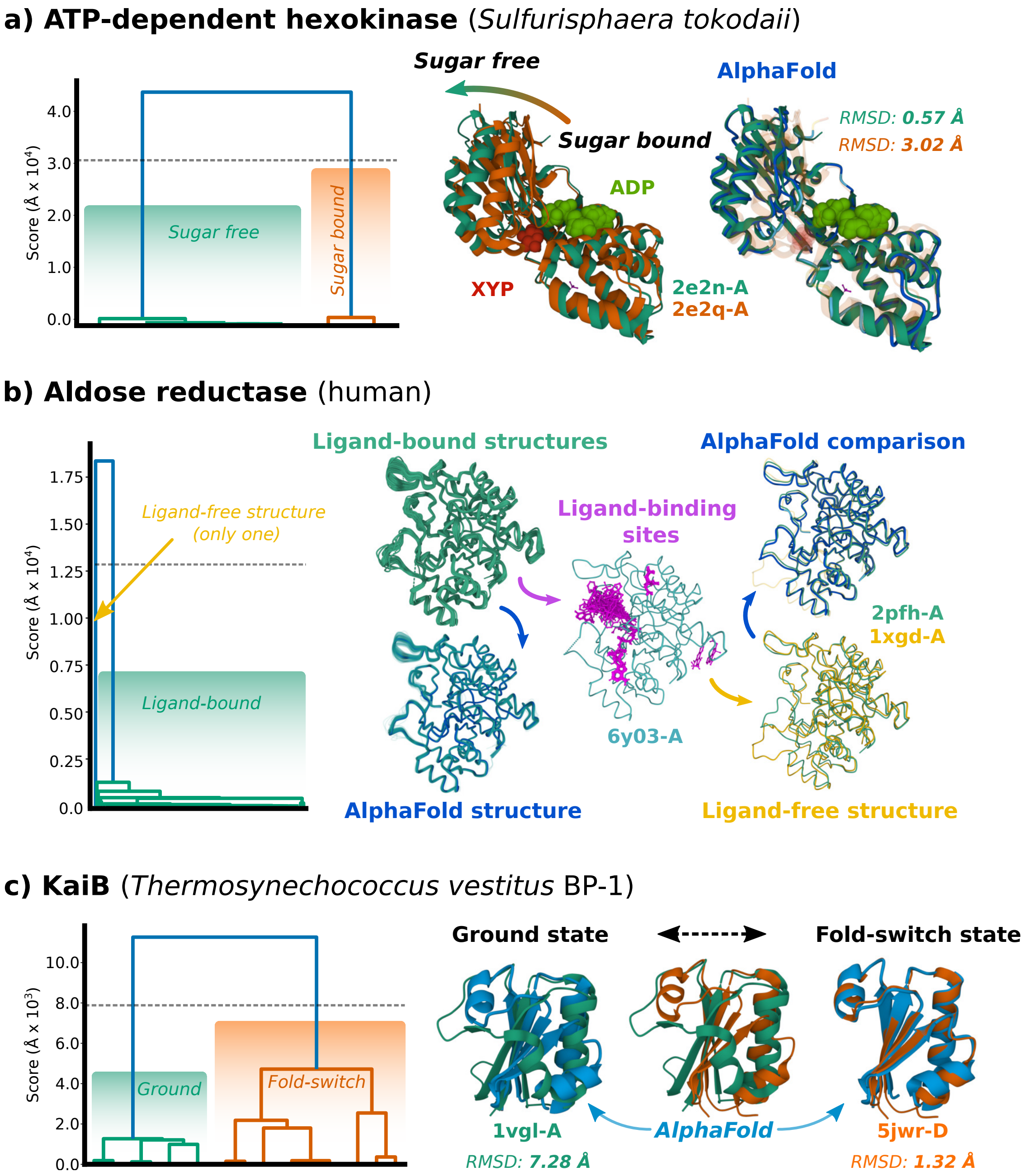

Figure 5: **Clustering results and superposed structures of biologically significant, recognised conformational states from the PDB archive.** Notable examples of clustering results from the PDB archive. UPGMA clustering dendrogram displayed on the left and Mol* figures of the superposed protein chains on the right. Chains are coloured by predicted conformational state and AlphaFold2 models are displayed in blue. RMSD displays the closeness in model coordinates between the predicted AlphaFold2 model and the experimentally determined structures from the PDB archive.

#### 3 Data and Code Availability

The entire pipeline is broadly separated into three components: 1) a segment definer, 2) the clustering application and 3) the structural aligner. Herein, the segment definer (§1.1) and clustering algorithm are described in detail (§1.2-1.5). Our segment definer also generates the appropriate input for the structural aligners GESAMT or SSM. The clustering application, named `protein-cluster-monomers` is open-source (available from the PDBe's GitHub repository) and can be deployed in separately from defined segments or structural alignment. The GitHub repository includes a README detailing how to run the application via a Python wrapper – allowing the program to be run locally on a set of parsed structures – or imported as a pip-installable package (`pip install protein-cluster-conformers`) into any Python >=3.10 scripts. The segment definer `protein-superpose` is written for the PDBe-KB's weekly release pipeline, but we are working to distribute an open-source version.

Accompanying the conformational state clustering pipeline is a benchmark dataset of peptide chains solved in either 'open' or 'closed' conformational states. The data set was compiled in order to check whether the GLOCON score (described in §1.4) captures a more even distribution of chains between conformations, compared to our previous use of the Q-score measurement returned by GSEAMT. The dataset was also used to check whether the GLOCON score could better separate true conformational states into clusters, which we found it did.
